# Impact of early life antibiotic and probiotic treatment on gut microbiome and resistome of very-low-birth-weight preterm infants

**DOI:** 10.1101/2024.12.18.629205

**Authors:** Raymond Kiu, Elizabeth M Darby, Cristina Alcon-Giner, Antia Acuna-Gonzalez, Anny Camargo, Lisa E Lamberte, Sarah Phillips, Kathleen Sim, Alexander G Shaw, Paul Clarke, Willem van Schaik, J Simon Kroll, Lindsay J Hall

**Affiliations:** Department of Microbes, Infection and Microbiomes, School of Infection, Inflammation and Immunology, College of Medicine and Health, University of Birmingham, Birmingham, UK; Institute of Microbiology and Infection, University of Birmingham, Birmingham, UK; Food, Microbiome and Health, Quadram Institute Bioscience, Norwich Research Park, Norwich, UK; Centro de Investigaciones en Microbiología y Biotecnología-UR (CIMBIUR), Facultad de Ciencias Naturales, Universidad del Rosario, Bogotá, Colombia; Health Sciences Faculty, Universidad de Boyacá, Tunja, Colombia; Faculty of Medicine, Imperial College London, London, UK; Norfolk and Norwich University Hospital, Norwich, UK; Norwich Medical School, University of East Anglia, Norwich, UK

## Abstract

Preterm infants (<37 weeks’ gestation) are often administered broad-spectrum antibiotics in hospitals due to their vulnerability to severe morbidity, including necrotising enterocolitis and sepsis. However, antibiotics can disrupt the development of early-life microbiota, potentially impairing gut immunity and colonisation resistance. Evidence shows that probiotics (e.g., certain *Bifidobacterium* strains) may help restore healthy gut microbiota. In this study, we examined the effects of probiotics and antibiotics on the preterm gut microbiome and resistome in two unique cohorts of 34 very-low-birth-weight, human-milk-fed preterm infants (moderate to very preterm), with one cohort receiving probiotics. Within each group, some infants were treated with antibiotics (benzylpenicillin and/or gentamicin) while others served as non-antibiotic treated controls. We performed shotgun metagenomic sequencing on 93 longitudinal faecal samples from 34 infants, generated >300 metagenome-assembled genomes, and obtained ∼90 isolate genomes through targeted culturomics, enabling analysis of the microbiome/resistome at species and strain levels. Additionally, we investigated *in vitro* horizontal gene transfer (HGT) capacity of preterm infant-derived multidrug-resistant (MDR) pathogen *Enterococcus* via neonatal gut models. Overall, probiotic supplementation significantly reduced antibiotic resistance gene prevalence, MDR pathogen load, and helped restore a typical early-life microbiota. However, the persistence of MDR pathogens like *Enterococcus*, with high HGT potential, highlights the need for ongoing surveillance in neonatal care. Our findings underscore the complex interactions between antibiotics, probiotics, and HGT in shaping the neonatal microbiome and support further research into probiotics for antimicrobial stewardship in preterm populations.

## Introduction

The World Health Organisation (WHO) estimates that over 10% of infants are born prematurely each year worldwide, defined as gestation of <37 weeks^1^. Among newborns, Very Low Birth Weight (VLBW) infants, those born weighing below 1500g, represent around 1.1-1.4%^2,3^. VLBW preterm infants have underdeveloped immune systems, making them particularly susceptible to morbidities such as Necrotising Enterocolitis (NEC)^4,5^ and sepsis^6,7^, often involving antibiotic-resistant bacteria. Due to these risks, preterm infants frequently reside in Neonatal Intensive Care Unit (NICUs) and are routinely administered broad-spectrum antibiotics, typically benzylpenicillin and gentamicin or derivatives, during their first days and weeks of life^8,9^. However, this early-life antibiotic exposure can disrupt the normal development of the gut microbiota^8,10,11^.

The use of antibiotics in preterm infants can also lead to an enrichment of antibiotic resistance genes (ARGs), collectively known as the gut resistome^12^. ARGs within gut bacterial communities can spread rapidly, primarily through horizontal gene transfer (HGT), occurring both within and between species, including from commensals to pathogens or vice versa, and this transfer often occurs via mobile genetic elements such as plasmids^13^.

Multidrug-resistant bacteria, including *Staphylococcus*, *Klebsiella*, *Enterococcus* and *Escherichia* are common in the preterm infant gut^5,8,14^, with their presence frequently linked to prolonged hospitalisation^15^, late-onset bloodstream infections^16^, and nosocomial infections in hospital settings^17–19^.

In response to these challenges, WHO has recommended probiotic supplementation for very preterm (<32 weeks’ gestation), human-milk-fed infants^20^. Probiotics, particularly *Bifidobacterium* spp. and *Lactobacillus* spp., are now increasingly used in NICUs^21^, with approximately 40% of NICUs in the UK adopting this practice^8,9^. Probiotic supplementation has been associated with reduced NEC incidence, lower mortality, fewer gut pathogens, and enhanced immune maturation^8,14,21–28^. Importantly, probiotics have also been observed to reduce the abundance of ARGs in the gut microbiota of preterm infants, bringing it closer to levels seen in full-term infants^29–32^.

To investigate the impact of probiotics on the preterm gut microbiome and resistome, we studied two cohorts of VLBW preterm infants, all exclusively fed human milk. Using shotgun metagenomics and genome-resolved approaches, we assessed microbiome species and strain dynamics and the effects of probiotics and antibiotics during the first three weeks of life. Additionally, a plasmid transfer experiment using *Enterococcus* in an infant gut model explored *ex vivo* ARG transfer.

## Results

We analysed the gut microbiome from 34 VLBW preterm infants (moderate to very preterm), divided into two main cohorts: the Probiotic-Supplemented (PS) cohort and the Non-Probiotic-Supplemented (NPS) cohort. Infants in the PS cohort received probiotics containing *B. bifidum* and *L. acidophilus*, while those in the NPS cohort did not receive any probiotics. These infants were selected as a subset from a larger study^8^. Within each cohort, some infants received empirical antibiotic treatment with benzylpenicillin and gentamicin, while others served as controls without antibiotic exposure (Fig. 1a). Faecal samples were collected weekly from preterm infants during the first three weeks of life, when possible (Table 1). These samples underwent processing, sequencing, and computational analysis to characterise the gut microbiome.

**Fig. 1.**
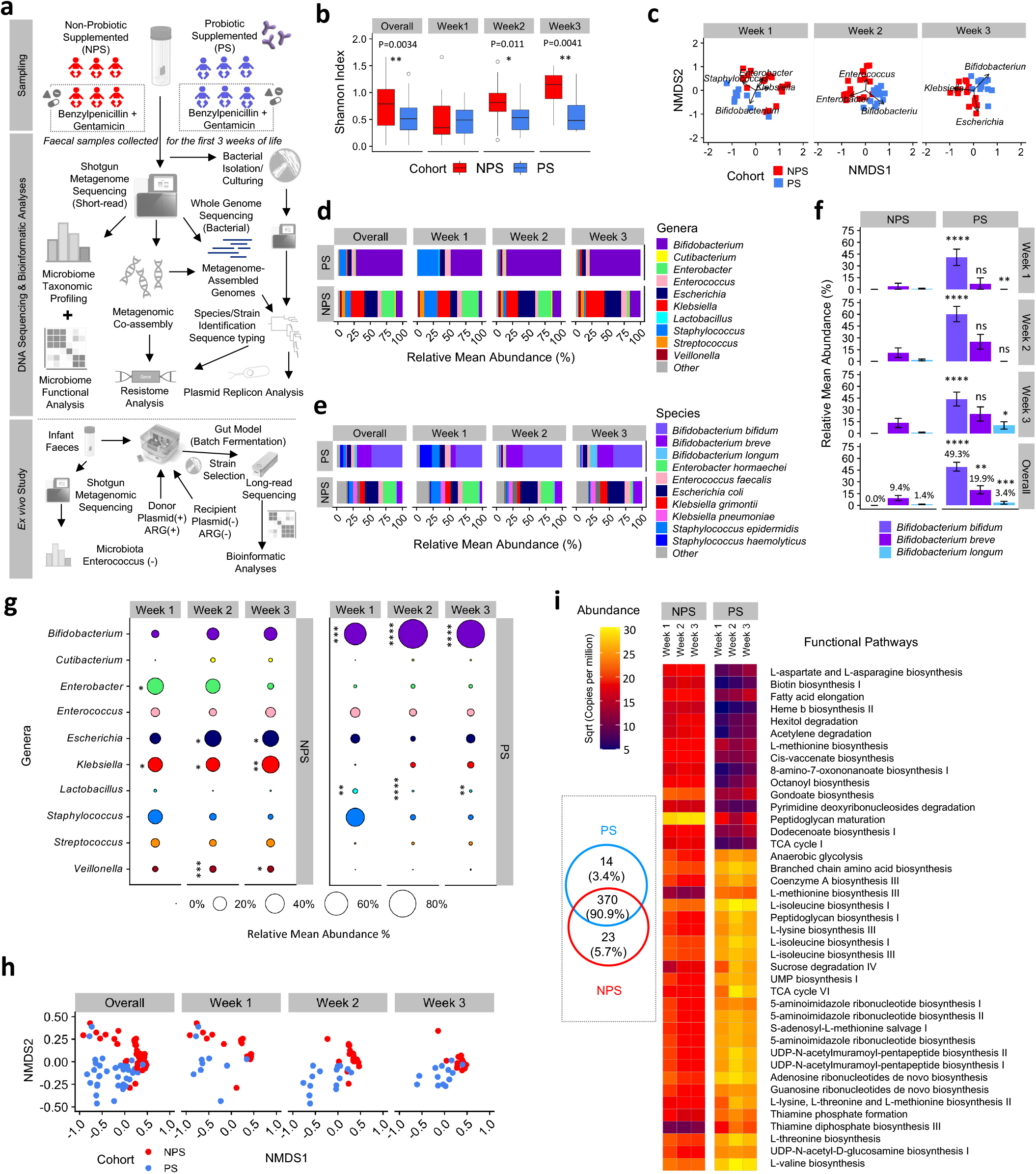
Neonatal-period preterm infant gut microbiome analysis and functional profiling. **a,** Schematic of this study with key details and relevant analyses. **b,** NMDS plots of 34 preterm infant gut metagenomes samples within the first 21 days of life (stratified by weeks and coloured by cohorts) and clustered based on genus-level Bray-Curtis dissimilarity. Significant taxa were labelled in the respective plots. **c,** Gut microbiome profiles of 34 infants in relative abundance (%) based on genus level data. Only top 10 most abundant genera across all samples were shown here. **d,** Gut microbiome profiles (as in **c**) of 34 infants based on species-level taxonomic assignment. Only top 10 most abundant bacterial species were shown here. **e,** Gut microbiome diversity indices of 34 infants in 2 cohorts, stratified by weeks (two-sided Wilcoxon test). **f,** Comparison of taxonomic abundance (%) of three key *Bifidobacterium* species across 2 cohorts (NPS vs PS) stratified by weeks. Percentages indicated are mean relative abundances in each species. Statistical significance was performed using two-sided Wilcoxon test (NPS vs PS with respect to weeks). * *P*<0.05, \*\**P*<0.01, \*\*\**P*<0.001, \*\*\*\**P*<0.0001. **g,** Mean taxa proportion of top 10 most abundant bacterial genera in both cohorts as shown in the bubble plots. The significance (two-sided Wilcoxon test) was compared NPS vs PS with respect to the time points (week). * *P*<0.05, \*\**P*<0.01, \*\*\**P*<0.001, \*\*\*\**P*<0.0001. **h,** NMDS plots of 34 preterm infant gut microbiome functional profiles stratified by weeks, NPS vs PS. **i,** Functional profiling heatmap showing 41 significant pathways profiled (alpha=0.05, LDA scores>2.0). The box-within-figure shows a Venn diagram of overlapping/unique functional pathways NPS vs PS based on a total of 407 pathways identified computationally.

**Table 1.** Summary of cohort characteristics: NPS cohort vs PS cohort. Cohort characteristics were compared using either Wilcoxon test, Student’s t-test or Fisher’s exact test where appropriate.

| Characteristic | NPS<br>(n=19) | PS<br>(n=15) |
| --- | --- | --- |
| **Demographics** |  |  |
| Male | 6 (32) | 10 (67) |
| Female | 13 (68) | 5 (33) |
| Median birth weight (IQR), g | 980 (1000-1325) | 1382* (1243-1451) |
| Median gestation at birth (IQR), weeks + days | 28+6 (28+2 - 30+1) | 30+0 (29+5 - 30+1) |
| Mode of delivery |  |  |
| Caesarean section | 17 (79) | 7 (47) |
| Normal vaginal delivery | 4 (21) | 8 (53) |
| **Antibiotic use in the first 3 weeks of life** |  |  |
| Median No. of days of treatment in treated infants (IQR) | 3.0 (3.0-5.0) | 3.0 (3.0-4.0) |
| No. of infants without treatment | 10 (53) | 3 (20) |
| No. of infants on short-course (1-3 day) treatment | 6 (32) | 8 (53) |
| No. of infants on long-course (>3 days) treatment | 3 (16) | 4 (27) |
| **Diet** |  |  |
| Breastmilk/donor breastmilk (week 1) | 19 (100) | 15 (100) |
| Breastmilk/donor breastmilk (week 2) | 19 (100) | 15 (100) |
| Breastmilk/donor breastmilk (week 3) | 18 (95) | 12 (80) |
| **Sample collection** |  |  |
| Median No. of samples collected (per infant) | 3.0 | 3.0 |
| Week 1 median time point, day (IQR), No. of infants (%) | 7.0 (7.0-7.8), 18 (95) | 6.0* (5.0-7.0), 11 (73) |
| Week 2 median time point, day (IQR), No. of infants (%) | 14.0 (14.0-14.0), 19 (100) | 13.0 (11.5-14.5), 15 (100.0) |
| Week 3 median time point, day (IQR), No. of infants (%) | 21.0 (20.8-21.3), 16 (84) | 20.0 (20.0-22.0), 13 (87) |
| **Sequencing depth** |  |  |
| Mean No. of sequencing reads | 10,578,191 | 10,869,232 |
| Mean metagenome size (bp) | 1,322,088,528 | 1,358,155,375 |

Data are presented as No. (%) unless otherwise specified. IQR, interquartile range. \* P<0.05.

### Probiotics support restoration of the early life gut microbiota

Gut microbiome diversity differed significantly between the NPS and PS cohorts, with the NPS cohort showing an increase in microbiome diversity during weeks 2 and 3, while the PS cohort maintained similar levels of diversity throughout (Fig. 1b). The overall gut microbiome profiles were markedly d istinct between the NPS and PS groups (Fig. 1c). The NPS infant gut microbiomes were characterised at the genus level by early-life pathobionts (Fig. 1d), including *Klebsiella*, *Enterobacter*, *Escherichia*, *Enterococcus*, and *Staphylococcus*, which on the species level were identified as *Klebsiella pneumoniae, Klebsiella grimontii*, *Enterobacter hormaechei*, *Escherichia coli*, *Staphylococcus epidermidis*, and *Staphylococcus haemolyticus* (Fig. 1e). In contrast, the gut microbiomes of PS infants were dominated by the genus *Bifidobacterium*, particularly *Bifidobacterium bifidum*, a major component of the Infloran probiotic provided to the infants, followed by *Bifidobacterium breve* and *Bifidobacterium longum* in weeks 2 and 3, reflecting the impact of probiotic supplementation (Fig. 1e). Notably, *B. breve* and *B. longum*, both associated with breastfeeding and recognised for promoting a healthy infant gut by breaking down complex carbohydrates, including human milk oligosaccharides (HMOs), were more abundant in PS infants compared to NPS infants (Fig. 1f).

Comparative analysis showed that *Bifidobacterium* was significantly more abundant in the PS cohort microbiota, while *Enterobacter*, *Escherichia*, and *Klebsiella* were more abundant in NPS infants (Fig. 1g). *Staphylococcus* was notably dominant only during the first week of life and declined in both cohorts by weeks 2 and 3, indicating its transient presence in the infant gut. Importantly, *Enterococcus faecalis*, a frequently multidrug-resistant bacterium, was also a prominent member of the gut microbiota in both cohorts.

Next, we examined the functional pathways within the gut microbiomes. Clustering of these pathways showed increasingly distinct patterns between the PS and NPS cohorts starting from week 2 (Fig. 1h). Across all samples, a total of 407 functional pathways were identified, with 84 pathways showing significantly different abundances between the two cohorts (Fig. 1i). Among these, 14 pathways (3.4%) were unique to the PS cohort, including sucrose degradation pathways, while 370 pathways (90.9%) were shared between both cohorts. In addition, 23 pathways (5.7%) were exclusively associated with NPS infants.

### Probiotic intervention is linked to suppression of ARGs in the gut resistome

We investigated the gut resistome in both NPS and PS cohorts, comprising infants treated empirically with benzylpenicillin and/or gentamicin, to assess the impact of these antibiotics on early gut microbiomes and resistomes over the first 3 weeks of life (Fig. 2a). Notably, the abundance of ARGs was significantly higher in NPS infants compared to PS infants across the first 3 weeks (Fig. 2b). Analysis of ARG diversity, defined by the number of antibiotic/drug classes represented, showed that NPS infant guts contained significantly more ARG types than those in the PS cohort, particularly in weeks 2 and 3 (Fig. 2c). Key ARG types in both cohorts included genes conferring resistance to aminoglycosides, macrolides- lincosamides-streptogramines (MLS), beta-lactamases, trimethoprim, and tetracycline, while ARGs conferring resistance to fluoroquinolones and colistin were exclusively found in NPS infants (Fig. 2d). Clustering analysis of resistome profiles revealed shared ARGs in both cohorts (48 ARGs), with 46 unique ARGs in NPS infants and 11 in PS infants, though the separation between groups was not clearly distinct (Fig. 1e, f).

**Fig. 2.**
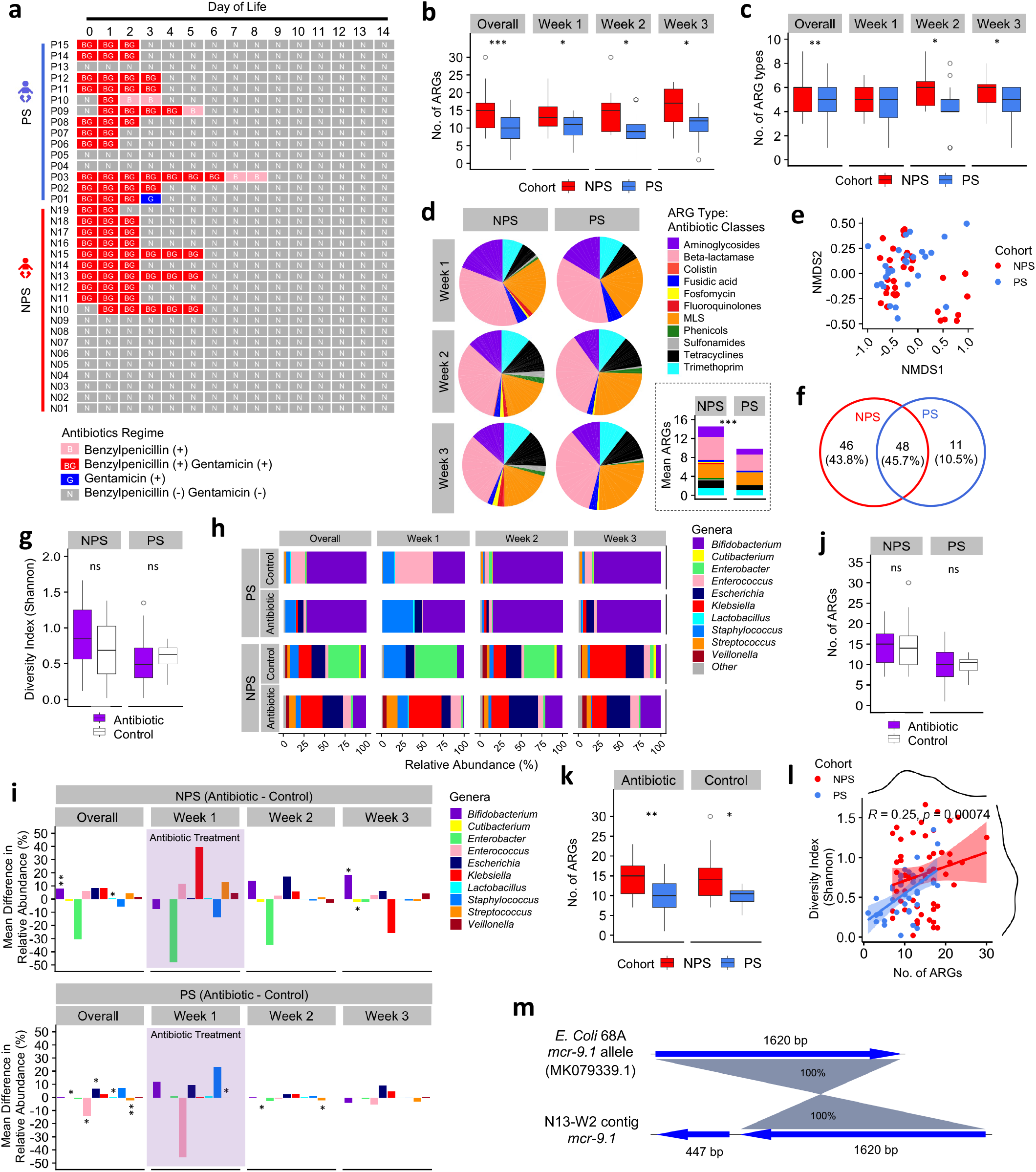
An in-depth profiling of neonatal preterm infant gut resistome. **a,** Schematic of antibiotics regime timeline in the preterm infants. **b,** Comparison of ARGs between NPS and PS cohorts. The significance was compared using two-sided Wilcoxon test. * *P*<0.05, \*\**P*<0.01, \*\*\**P*<0.001. **c,** Number of ARG types (antibiotic/drug classes against which encoding ARGs are resistant) comparison between NPS and PS cohorts stratified by time points. Statistical analysis was performed using two-sided Wilcoxon test. * *P*<0.05, \*\**P*<0.01. **d,** Proportions of ARG types in both NPS and PS cohorts stratified by weeks. The mini bar chart within sub-figure represents the mean number of ARGs in both NPS and PS cohorts coloured by ARG types. Statistical analysis was performed using two-sided Wilcoxon test. \*\*\**P*<0.001. **e,** NMDS plot of ARG profiles of individuals in bot NPS and PS cohorts. **f,** Venn diagram showing number/percentages of unique and sharing ARGs between NPS and PS cohorts. **g,** Shannon index (diversity index) comparison between Antibiotic vs Control (untreated) in NPS and PS cohorts. **h,** Preterm gut microbiome dynamics (Top 10 genera were shown) in the first three weeks of life, stratified by antibiotic treated individuals and control group (without antibiotic treatment). **i,** Increase/reduction of taxonomic abundance (Top 10 genera) in Antibiotic group (vs Control) in both NPS and PS cohorts. Shaded area represents antibiotic treatment period, infants in Antibiotic group were treated with antibiotic only in their first week of life (3 median days). Statistical analysis was performed using two-sided Wilcoxon test. . * *P*<0.05, \*\**P*<0.01. **j,** No. of ARGs comparison between Antibiotic vs Control (untreated) in both NPS and PS cohorts. **k,** Number of ARGs was compared NPS vs PS, stratified by antibiotic usage. Statistical analysis was performed using two-sided Wilcoxon test. * *P*<0.05, \*\**P*<0.01. **l,** Correlation plot of (Shannon) diversity index vs ARGs using all available data points. Scatter plot was coloured by cohorts accordingly. *R* represents Kendall rank correlation coefficient. **m,** Genomic mapping of a computationally extracted sequence contig with *mcr-9.1* ARG gene against a *mcr-9.1* ARG from *E. coli* (NCBI nucleotide database) with 100% matching nucleotide identity.

To examine the impact of empirical antibiotic therapy on the gut microbiome, we compared Antibiotic-treated vs Control (non-antibiotic-treated) groups within each cohort. Overall, no significant difference in microbiome diversity was observed between Antibiotic and Control groups (Fig. 2g). In the NPS cohort, the Antibiotic group showed higher gut microbiome diversity from the first week, which continued through weeks 2 and 3, while the Control group’s microbiome diversity increased in later weeks; both groups were dominated by key pathobionts (Fig. 2h). In the PS cohort, apart from an initial increase in *Staphylococcus* in Antibiotic-treated infants and *Enterococcus* in Controls, *Bifidobacterium* dominated in weeks 2 and 3. Taxonomic abundance analysis revealed no significant overall differences in abundance for Antibiotic vs Control in the NPS cohort, except for *Bifidobacterium* and *Lactobacillus* (Fig. 2i, top). In the PS cohort, Antibiotic-treated infants had significantly higher levels of *Cutibacterium*, *Escherichia*, and *Lactobacillus*, with reduced *Enterococcus* and *Streptococcus* over 3 weeks (Fig. 2i, bottom).

To further understand antibiotic impacts on the resistome, we compared ARG profiles in Antibiotic vs Control groups within each cohort. No significant differences were observed in ARG counts between these groups in either cohort (Fig. 2j). However, both Antibiotic-treated and Control infants in the NPS cohort had higher ARG counts than their counterparts in the PS cohort, suggesting a potential ARG-suppressive effect of *Bifidobacterium bifidum* in the preterm gut (Fig. 2k).

We also assessed whether microbiome diversity correlated with ARG abundance in preterm infants. Statistical testing indicated a weak, but statistically significant, positive correlation between increased microbiome diversity and ARG levels (correlation coefficient R=0.25; P=0.00074; Fig. 2l). Notably, a colistin resistance gene, *mcr-9.1*, was detected with 100% nucleotide identity in the gut microbiome of a preterm infant around 2011-2012 (Fig. 2m), predating its identification as a resistance gene in 2019, though it could not be computationally mapped to a specific bacterial strain in this study.

### Strain-level gut-derived pathobiont resistome analysis

To examine gut resistomes at the strain level, we applied genome-resolved metagenomics, recovering 322 high-quality metagenome-assembled genomes (MAGs) from shotgun metagenome sequences and incorporating 89 novel isolate genomes cultured from preterm infant faecal samples. Together, these genomes represent 27 genera and 47 species.

Resistance-gene searches and statistical analyses were conducted on these bacterial genomes to characterise the early-life gut resistomes.

Following dereplication of all genomes, including MAGs, at a strain-level cut-off of 99.9% ANI, we obtained 195 representative genomes for further resistome analysis (Fig. 3a). In total, 10 ARG types were identified across bacterial strains (Fig. 3a), with *Enterococcus*, *Escherichia*, *Klebsiella*, and *Staphylococcus* identified as the top four resistant genera, each exhibiting a significantly higher ARG count per strain vs the rest (Fig. 3b). Further examination of these genera showed a trend towards lower ARG counts in *Enterococcus*, *Escherichia*, and *Klebsiella* strains within the PS cohort compared to NPS, though differences were not statistically significant (Fig. 3c).

**Fig. 3.**
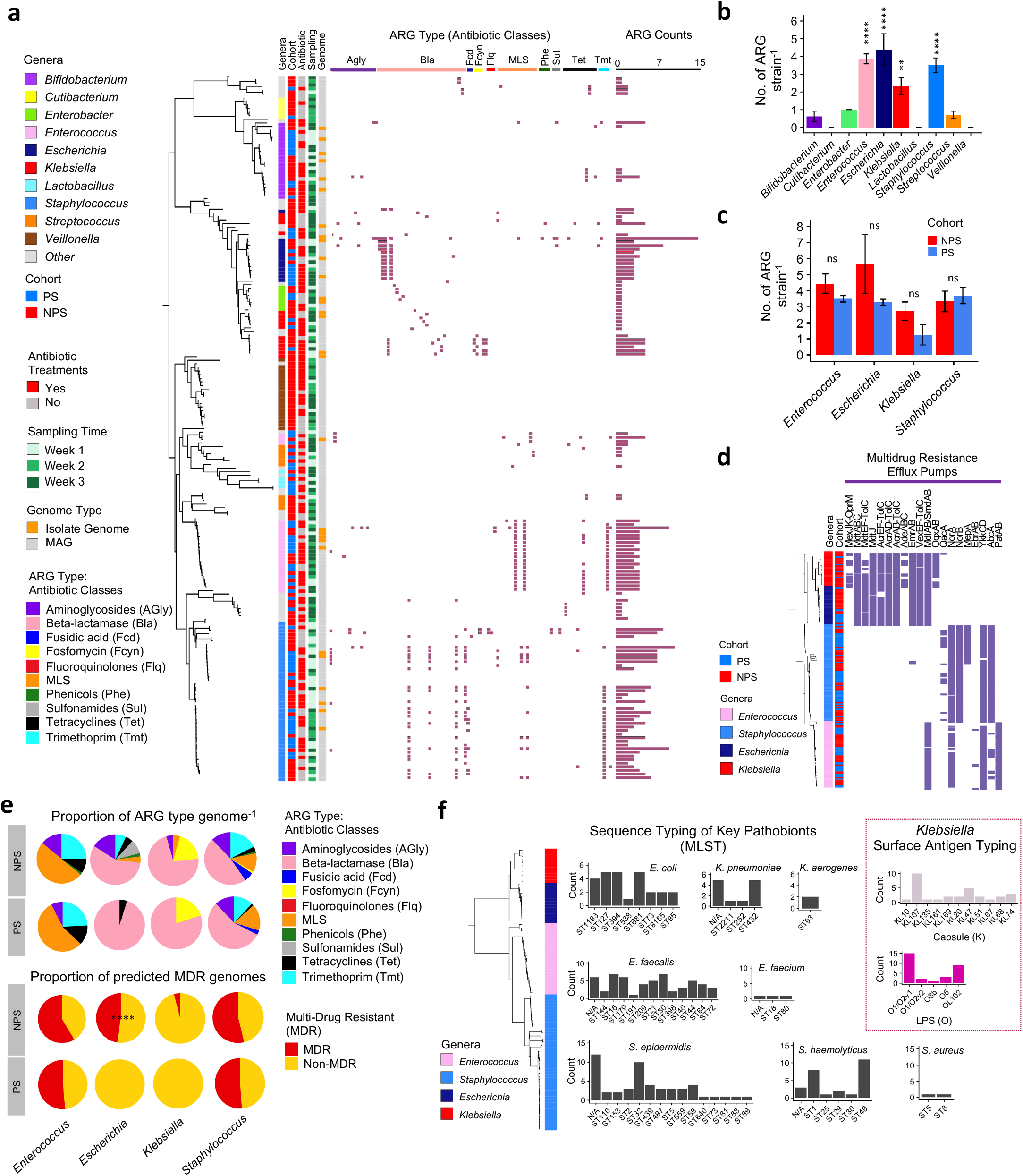
Strain-level gut-derived pathobiont resistome analysis. **a,** Strain-level neighbor-joining tree of bacterial strains (n=195) aligned with clinical metadata, ARG profiles and gene counts, including both isolate genomes and metagenome-assembled genomes (MAGs) derived from a total of 411 individual bacterial genomes. **b,** ARG count per strain comparison across Top 10 genera. Statistical analysis were performed using Kruskal Wallis test and post-hoc Dunn’s test (FDR adjusted). Significance was compared against *Bifidobacterium*. \*\**P*<0.01, **** *P*<0.0001. **c,** ARG count per strain comparison between Top 4 drug-resistant pathobiont genera including *Enterococcus*, *Escherichia*, *Klebsiella* and *Staphylococcus*. Statistical analysis between cohorts were performed using two-sided Wilcoxon test. ns: non-significant. **d,** Multidrug resistance efflux pump profiles (n=20) of Top 4 drug-resistant pathobiont genera *Enterococcus*, *Escherichia*, *Klebsiella* and *Staphylococcus* genomes (n=205). **e,** (Top) Proportion of ARG type per genome (n=205) in both NPS and PS cohorts. (Bottom) Proportion of MDR/non-MDR genomes in both NPS and PS cohorts. MDR is defined as having three or more predicted ARG types (resistant against drugs of three or more antibiotic classes). Statistical test was performed using Fisher Exact test. \*\*\*\**P*<0.0001. **f,** Multi-locus Sequence Typing (MLST) analysis of key pathobiont genomes (n=193) including Top 4 pathobiont genera *Enterococcus*, *Staphylococcus*, *Escherichia* and *Klebsiella*. Additional surface antigen typing (Kaptive) for *Klebsiella* spp as shown in the box-within-figure.

A deeper analysis of multidrug resistance (MDR) efflux pumps across these resistant genera revealed multiple efflux pumps, with *Escherichia* and *Klebsiella* strains encoding the highest number (n=13), followed by *Staphylococcus* (n=7) and *Enterococcus* (n=6) (Fig. 3d). ARG type analysis further demonstrated the widespread presence of beta-lactamase resistance genes in *Escherichia*, *Klebsiella*, and *Staphylococcus* strains, with aminoglycoside resistance genes prevalent in NPS *Enterococcus* strains (Fig. 3e, top). Analysis of MDR capacity showed that *Enterococcus* ranked highest among these resistant genera, with over half of its strains harbouring ARGs conferring resistance to more than three antibiotic classes, followed by *Staphylococcus*. Notably, none of the PS-associated *Klebsiella* (n=5) or *Escherichia* (n=12) strains were MDR, while 47.6% of NPS-associated *Escherichia* (n=31) strains exhibited MDR characteristics (Fig. 3e, bottom).

To assess clinical relevance, we sequence-typed a total of 193 key resistant pathobiont genomes from *Escherichia*, *Klebsiella*, *Enterococcus*, and *Staphylococcus* (Fig. 3f) using the multi-locus sequence typing (MLST)^33^ scheme. Dominant sequence types (STs) included ST1193, ST127, ST394, and ST681 for *E. coli*, while ST432 was prominent for *K. pneumoniae*. In *Enterococcus*, 12 STs were identified, including six potential novel STs. *S. epidermidis* displayed 14 STs, with ST32 in 10 genomes and 12 potential novel STs. Of particular note, key *S. haemolyticus* STs included methicillin-resistant ST1, linked to NICUs previously^34^ and ST49, associated with children and often resistant to methicillin^35,36^. We also performed surface antigen typing for *Klebsiella*, identifying KL10 (n=10) as the most common capsule (K) type and O1/O2v1 (n=15) as the most prevalent LPS (O) type in the *Klebsiella* genomes (Fig. 3f, right).

### Strain-level mobilome analysis and circulation of strains within NICUs

To investigate whether plasmid carriage is linked to the resistome, we performed a plasmid replicon search across 195 strain-level bacterial genomes (Fig. 4a). Comparison between PS and NPS cohorts revealed significantly higher plasmid counts in the NPS cohort (Fig. 4b). To further assess the role of antibiotics in the mobilome, we compared plasmid counts between Antibiotic-treated and Control groups within both cohorts. Although Antibiotic-treated strains showed a higher plasmid count, the difference was not statistically significant in either cohort (Fig. 4c) or in the overall comparison (Fig. 4d).

**Fig. 4.**
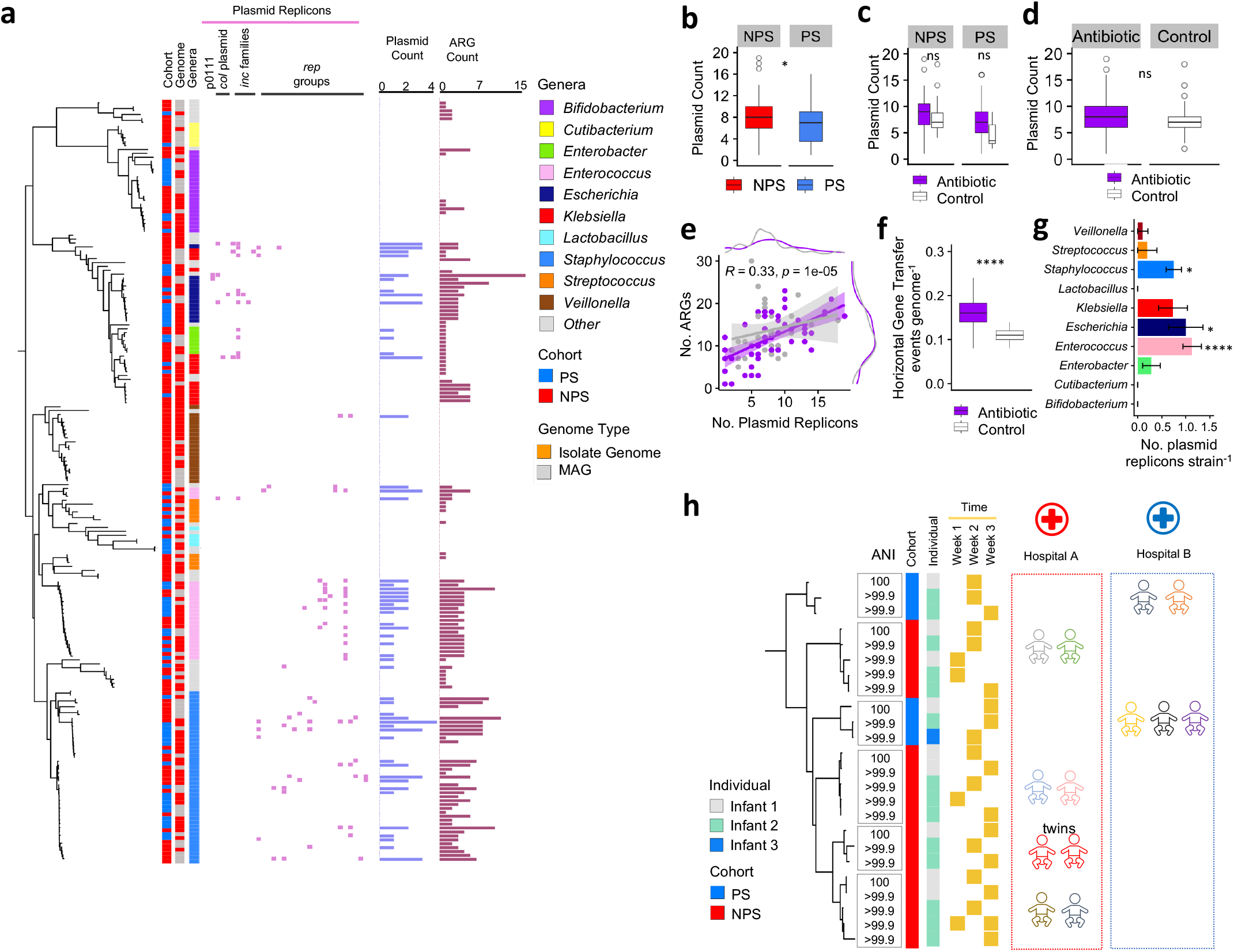
Strain-level mobilome analysis and persistence of strains in preterm infant gut microbiome. **a,** Plasmid replicon profiles of preterm infant gut-associated bacterial strains (n=195) aligned with a strain-level neighbor-joining tree. Plasmid and ARG counts were also included for comparison. **b,** Gut microbiome plasmid count comparison NPS vs PS. Statistical test was performed using Wilcoxon test. \**P*<0.05. **c,** Gut microbiome plasmid count comparison Antibiotic vs Control, stratified by cohort NPS vs PS. Statistical test was performed using Wilcoxon test. ns: non-significant. **d,** Gut microbiome plasmid count comparison between Antibiotic vs Control in all preterm infant metagenome samples. **e,** Correlation plot of ARGs vs plasmid count (in all preterm infant metagenome samples) using all available data points. Scatter plot was coloured by Antibiotic usage accordingly, where purple represents infants administered with antibiotic while white indicates untreated infants. *R* represents Kendall rank correlation coefficient. **f**, HGT events per genome (n=411) between Antibiotic groups and Control. Statistical analysis was performed using Wilcoxon test. \*\*\*\**P*<0.0001. **g,** Plasmid replicon count per strain comparison across the Top 10 genera. Statistical analysis were performed using Kruskal Wallis test and post-hoc Dunn’s test (FDR adjusted). Significance was compared against *Bifidobacterium*. \**P*<0.05, **** *P*<0.0001. **h,** Potential dissemination of *Enterococcus* strains among certain preterm infants in the NICUs. Strain-level cut-off ANI>99.9%.

Next, we examined the correlation between plasmid replicon count and ARG count, finding a weak positive correlation (R=0.33; P=0.00001; Fig. 4e). Notably, the Antibiotic-treated group exhibited a significantly higher frequency of potential horizontal gene transfer events compared to the Control group (Fig. 4f). Among key pathobiont genera, *Enterococcus*, *Escherichia*, and *Staphylococcus* were identified as top plasmid carriers, with *Enterococcus* ranking highest in both ARG and plasmid carriage (Fig. 4g). Given its prominence as an MDR pathogen and its relative understudy in preterm infants, we focused on tracking *Enterococcus* transmission among infants in both cohorts.

Average nucleotide identity (ANI) analysis at a strain-level cut-off of 99.9% indicated 6 *Enterococcus* strains were circulating among infants, with four pairs of infants (unrelated) in Hospital A (one of which was a twin) and one pair in Hospital B carrying identical strains (Fig. 4h). Additionally, in Hospital B, three other unrelated infants were colonised with the same *Enterococcus* strain, underscoring the potential for nosocomial transmission of this MDR pathogen.

### AMR plasmid transfer between Enterococcus strains occurs within an infant gut model

Given the clinical significance of *Enterococcus faecium* as a multidrug-resistant (MDR) pathogen and its relatively limited study in infant microbiomes, we hypothesised that ARG-encoding plasmids could be transferred between *E. faecium* strains within the infant gut. To test this, we used two donor strains, both originated from preterm infants, and a plasmid-free, gentamicin-sensitive recipient strain (64/3) in a plasmid transfer experiment (Fig. 5a). We utilised a gut model using *Enterococcus*-free infant faecal slurry as the culture medium to simulate the neonatal gut environment (Fig. 4b). Following 24 hours incubation, we isolated *Enterococcus* colonies on selective media and performed WGS. Genome analysis revealed the successful transfer of a ∼137kb mega-plasmid encoding the aminoglycoside resistance gene *aac6-aph2* to plasmid-free recipient strain 64/3 (transformants D1 and D3), thereby conferring gentamicin resistance to these strains (Fig. 5b, 5c and 5d).

**Fig. 5.**
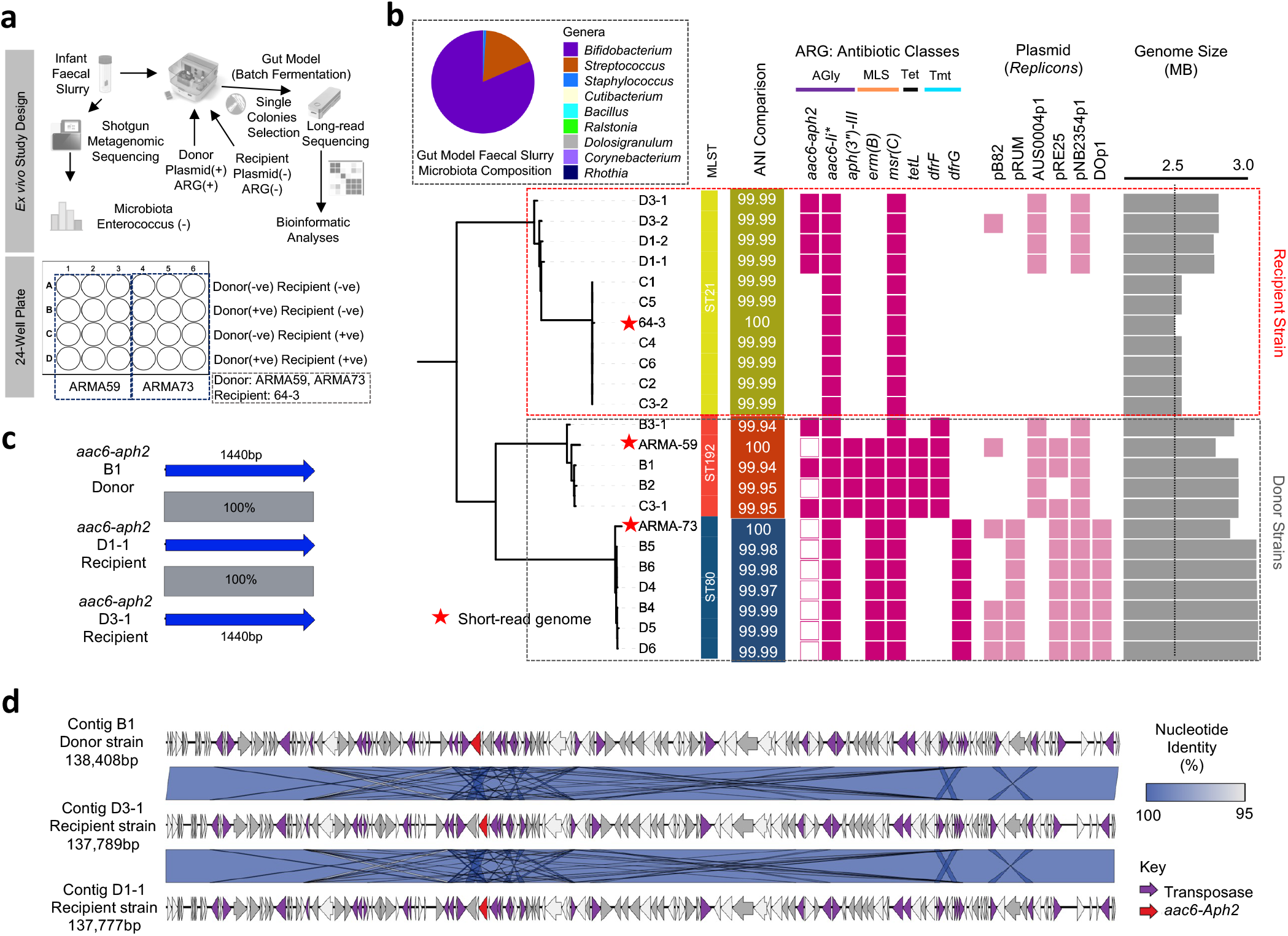
Gentamicin-resistant plasmid transfer study in *Enterococcus.* **a,** Schematic of *in vitro* study workflow and relevant analyses with the study design in 24-well plate template. **b,** Antibiotic resistance gene profiling of donor and recipient strains, colonies singly picked from selective agars supplemented with appropriate antibiotics and whole-genome sequenced via long-read/short-read sequencing platform. A neighbour-joining tree was constructed to reflect the relatedness between isolate genomes, aligned with plasmid replicon predicted in the genomes in tandem with calculated genome sizes. The box-within-box (top left) was shotgun metagenome contents (microbial taxa) of infant faecal slurry that was used in the gut model experiment. \**aac6-Ii* is known as a chromosomal-encoded aminoglycoside acetyltransferase in *Enterococcus* spp. **c,** Comparison of computationally extracted antibiotic resistance gene *aac-aph2* from representative donor strain B1, recipient (transformant) strains D1-1 and D3-1. **d,** Comparison of complete contig sequences that encoded gentamicin resistance gene *aac6- aph2,* predicted to be either whole or part of potential mobile genetic elements including plasmids.

## Discussion

In this study, we used genome-resolved metagenomics to analyse the resistome and microbiome of preterm infants during their first 3 weeks of life in neonatal units, highlighting the complex effects of antibiotics, probiotics, and HGT on the gut microbiome. Our findings suggest that probiotic supplementation with *B. bifidum* and *L. acidophilus* not only supports beneficial microbial communities but also plays a role in reducing MDR bacteria and overall ARG carriage.

One of the strengths of this study is the inclusion of both PS and NPS infants who did not receive antibiotics, which is uncommon considering the high antibiotic prescription rates among these at-risk patients (55-87% for very to extremely preterm infants in the UK)^37^, allowing us to begin disentangling the specific effects of probiotics and antibiotics on the preterm gut microbiome and resistome. This carefully selected sub-cohort (n=34) from a larger observational study (n=234) consists exclusively of preterm infants (<33 weeks’ gestation) who were matched for diet and birth mode, providing a relatively uniform baseline^8^. Moreover, in cases where empirical antibiotic therapy was administered, the duration and types of antibiotics (benzylpenicillin and/or gentamicin, the most frequently prescribed antibiotics in UK NICUs^37^) were strictly limited and standardised to less than 1 week. Despite this exposure, we observed minimal impact on overall microbiome diversity within both PS and NPS cohorts, suggesting that early-life antibiotic exposure over a short period may not have an immediate effect on preterm microbiome diversity. However, it is possible that the limited 3-week sampling window did not capture potential longer-term shifts induced by antibiotic exposure. This observation aligns with previous studies that reported minimal immediate impact of antibiotics on early microbiome composition, while indicating that more pronounced changes may emerge over extended periods^38^.

However, the administration of antibiotics, though limited to an average duration of median 3 days, did specifically impact microbiome composition by promoting the proliferation of certain genera with pathogenic potential, including *Klebsiella* and *Enterococcus*, in the first week, with some of these strains persisting into the second week. *Enterobacter*, a genus less associated with multiple ARGs in this study, was observed to decline in the NPS cohort after antibiotic exposure, indicating potential susceptibility to the antibiotic regimen. In contrast, *Bifidobacterium* - particularly in the PS cohort - appeared to exert a protective effect, as evidenced by a marked reduction in *Enterococcus* abundance. This decrease could potentially be attributed to the carbon source depletion promoted by *Bifidobacterium* specialised HMO utilisation in human milk-fed gut environment, and reduced niche pH, which may inhibit pathobiont colonisation and growth^39,40^.

Overall, we identified a diverse range of ARGs in the gut microbiomes of preterm infants, with key ARG classes including genes encoding resistance to aminoglycosides, macrolides-lincosamides-streptogramins (MLS), beta-lactamases, trimethoprim, and tetracyclines. These ARG classes are clinically relevant in NICUs, where aminoglycosides and beta-lactams, e.g. gentamicin and benzylpenicillin, are frequently empirically administered from birth to cover possible infection^8,37^. The presence of these ARGs, underscores the potential for reduced antibiotic efficacy in this vulnerable population and highlights the need for careful and improved antibiotic stewardship in NICU settings. Efflux pumps, which facilitate multidrug resistance by expelling a broad range of antibiotics^41^, were also prevalent across multiple strains, particularly within *Escherichia* and *Klebsiella*, further complicating treatment options for infections by these pathogens in preterm infants. Notably, we detected an *mcr-9.1* gene, conferring resistance to colistin - a last-resort antibiotic -within the gut microbiome of a NPS infant. This gene was identical to a homolog found in *E. coli* strain 68A though it could not be definitively linked to a specific bacterial species in our cohort due to limitations in short-read sequencing data, as it appeared only as a single contig (Fig. 2m). The *mcr-9.1* gene was first identified in *Salmonella enterica* in 2019^42^, where it conferred phenotypic resistance to colistin, and has since been associated with *Enterobacter* and *Klebsiella* species, often on mobile IncHI2 plasmids^43^. The presence of *mcr-9.1* in our cohort raises concerns about the silent circulation of colistin resistance genes within the gut microbiome of preterm infants, which may complicate future treatment options, particularly in cases where conventional antibiotics fail.

We also observed an increase in ARG abundance over time in NPS cohort, indicating that ARG acquisition may be driven by the introduction of new bacteria, likely hospital-acquired, rather than ARG evolution within the closed ecological system of the infant gut (at least across the short sampling widow). This is further supported by our strain-level analyses, where we observed the persistence of certain pathobionts, such as *Enterococcus*, across multiple time points. This persistence, along with the detection of shared strains among infants within the same hospital, suggests a high likelihood of nosocomial transmission of MDR bacteria. These findings align with reports from other NICU studies, where *Enterococcus*, *Escherichia*, and *Klebsiella* are common nosocomial pathogens and are often implicated in late-onset infections in preterm infants^32,44,45^. Indeed, we identified specific STs linked to neonatal infections, such as *E. coli* ST1193^46^, ST73 and ST95^46,47^, and *K. pneumoniae* ST432^48^ strains, which are typically drug-resistant strains and of clinical significance. *Enterococcus*, frequently identified as a prominent ARG carrier in our cohort, exhibited multiple STs, including novel ones not previously reported, and certain strains were shared across different individual infants within the same hospital. This sharing among unrelated infants further supports the notion of nosocomial transmission of *Enterococcus*, a common cause of late-onset infections in NICUs. Thus, the frequent colonisation with hospital-acquired, ARG-bearing bacteria in the absence of probiotics underscores the importance of continuous surveillance and infection control in NICUs.

Our findings on the high prevalence of strains carrying plasmids, and in particular MDR plasmids, is particularly concerning, as it indicates that these strains could serve as reservoirs for resistance genes that may spread to other non-MDR strains within the same species as well as other genera via HGT. Notably, our findings indicate that antibiotic treatment promotes a higher prevalence of LGT events within the neonatal gut, likely due to the selective pressure antibiotics place on microbial populations. This pressure appears to favour the retention and spread of mobile genetic elements carrying ARGs, increasing HGT dynamics in antibiotic-treated infants. Indeed, our plasmid transfer experiment within *E. faecium* further emphasises the significance of plasmid-mediated ARG transfer in the neonatal gut as we demonstrated the successful transfer of a ∼137kb mega-plasmid carrying an aminoglycoside resistance gene (*aac6-aph2*) to a plasmid-free, gentamicin-sensitive recipient strain, conferring phenotypic gentamicin resistance. It is important to note that our genomic analysis indicated only a weak correlation between overall ARG abundance and plasmid replicon count, suggesting that plasmids are not the sole contributors to ARG dissemination, and alternative routes of HGT are likely driving ARG spread^49^.

The high-resolution sequencing data in this study underscored the beneficial impact of probiotic supplementation on microbiome development in preterm infants. PS infants showed an earlier and more robust colonisation of infant-associated *Bifidobacterium* species, particularly *B. breve* and *B. longum* subsp. *infantis*, compared to NPS infants^50–52^. The probiotic *B. bifidum* likely facilitated the establishment of these beneficial species through cross-feeding on HMOs^53,54^, promoting a microbiome composition more similar to that of term infants. Indeed, this early and abundant presence of multiple *Bifidobacterium* species appeared to enhance pathogen colonisation resistance significantly, as evidenced by the limited presence of pathobionts, such as *Klebsiella* and *E. coli*, within the PS cohort.

Consistent with previous studies, our results also showed that PS infants harboured significantly fewer ARGs than NPS infants^29^. *Bifidobacterium* supplementation has been associated with lower ARG abundance in the infant gut microbiome, likely due to the competitive exclusion of ARG-rich pathobionts^55,56^. In our study, the higher abundance of *Bifidobacterium* in the PS cohort correlated with a reduced ARG load, particularly in genera commonly associated with MDR pathogens, such as *Enterococcus*, *Escherichia*, and *Klebsiella*. Notably, *Escherichia* and *Klebsiella* genomes in PS infants did not exhibit MDR characteristics, highlighting the potential for probiotic supplementation to mitigate MDR pathogen prevalence in the preterm gut. However, while plasmid analysis revealed a greater number of plasmids in the NPS cohort compared to PS infants, suggesting that probiotics may help reduce the plasmid pool carrying ARGs and, by extension, limit the ARG reservoir, we still observed plasmid transfer *in vitro* even in the presence of high levels of *Bifidobacterium*, suggesting that presence of beneficial bacteria may not fully inhibit HGT. This observation underscores a critical point: while probiotics such as *Bifidobacterium* support beneficial microbiome composition and may suppress ARG-rich pathobionts, they do not necessarily prevent HGT or plasmid transfer, particularly in the antibiotic-exposed gut where HGT may be more common. This emphasises the need for further studies to evaluate the role of probiotics not only in microbial colonisation but also in their potential impact on HGT dynamics, particularly in environments where antibiotics are heavily used.

Finally, one limitation of our study was the relatively short sampling period, which cannot capture longer-term impacts of antibiotics and probiotics on microbiome development.

Additionally, the study involved samples from two separate hospital sites, which allowed us to examine baseline and probiotic-supplemented conditions but inevitably introduces potential confounders from local variability in practices. Future studies involving larger sample sizes, extended sampling windows, and multi-site designs could provide more comprehensive insights into the dynamic interactions between antibiotics, probiotics, and the neonatal microbiome.

In conclusion, our investigation provides a comprehensive overview of the preterm gut resistome and microbiome, demonstrating the role of probiotic supplementation in reducing ARG prevalence and pathogen load. However, the persistence of MDR pathogens such as *Enterococcus*, with its high plasmid carriage and demonstrated capacity for ARG transfer, even in a *Bifidobacterium*-dominant ecosystem, underscores the need for continued surveillance and targeted intervention strategies in NICUs to minimise the risk of colonisation and subsequent infections by MDR bacteria. Our findings highlight the complex interplay between antibiotics, probiotics, and HGT in shaping the neonatal microbiome, and provide a platform for further research into the role of probiotics in antimicrobial stewardship and infection control in vulnerable preterm populations.

## Methods

### Cohort and sample selection

A sub-set of 93 samples in total were selected from a previously published observational cohort study^8^. All NICUs in this study applied similar protocols for feeding, prescription of antibiotics and the use of prophylactic antifungal drugs. Hospital for PS cohort routinely prescribed all VLBW infants with oral probiotic supplementation (Infloran®, Desma Healthcare, Switzerland) twice daily (from birth until ∼34 weeks post-menstrual age), while infants in NPS cohort were not supplemented with probiotics. Probiotic supplements contained *Bifidobacterium bifidum* (1 x 10^9^ cfu) and *Lactobacillus acidophilus* (1 x 10^9^ cfu). Samples from both cohorts (NPS and PS) were selected for this study based on the following criteria: VLBW preterm infants with gestational age < 34 weeks, infants were solely given antibiotics benzylpenicillin and/or gentamicin, and age-matched infants without antibiotics, infants were all fed with breastmilk and/or donor breastmilk in both cohorts. A total of 34 VLBW preterm infants were selected for this study. Longitudinally collected faecal samples at week 1, week 2 and week 3 (first 3 weeks of their NICU stay) were recruited into this study (Supplementary Table 1).

### Cohort characteristics

This study is a ‘controlled’ sub-study from the original observational longitudinal study^8^ as published previously. Summary of the cohort characteristics are presented in **Table 1**. Faecal slurry for the plasmid transfer experiment was originally from five infants recruited in PEARL study^57^.

### Ethical approval

Faecal sample collection from Norfolk and Norwich University Hospital (BAMBI study) was approved by the Faculty of Medical and Health Sciences Ethics Committee at the University of East Anglia (UEA), and followed protocols laid out by the UEA Biorepository (License no: 11208). Faecal sample collection Imperial Healthcare NICUs (NeoM study) was approved by West London Research Ethics Committee (REC) under the REC approval reference number 10/H0711/39. In all cases, medical doctors and nurses recruited infants after parents gave written consent. The PEARL study has been reviewed and agreed by the Human Research Governance Committee at the Quadram Institute Bioscience and the London-Dulwich Research Ethics Committee (reference 18/LO/1703) and received written ethical approval by the Human Research Authority. IRAS project ID number 241880^57^.

### Genomic DNA extraction and Shotgun Metagenomic Sequencing

FastDNA Spin Kit for Soil (MP Biomedicals) was utilised to extract genomic DNA from infant faeces following manufacturer instructions, with an extended 3 min bead-beating on FastPrep tissue homogeniser (MP Biomedicals), additionally DNA was eluted with 55°C sterile pure water. Genomic DNA concentration quantification followed using a Qubit 2.0 fluorometer (Invitrogen). DNA samples were then subject to established Illumina paired-end library preparation prior to Shotgun Metagenomic Sequencing on Illumina HiSeq 2500 to generate 125-bp paired-end reads (FASTQ), which was performed at the Wellcome Trust Sanger Institute (Supplementary Table 2).

### Bacterial isolation work, DNA extraction

A targeted bacterial isolation work was carried out to isolate the most abundant taxa present in the preterm infant faecal samples – *Bifidobacterium*, *Enterococcus*, *Staphylococcus*, *Klebsiella* and *Escherichia* genera. Briefly, 25-50 mg of faecal samples were homogenised in 5 ml of phosphate buffer saline (PBS) by vortexing. Homogenates were then serially diluted to 10^-4^ in PBS and aliquots of 100 ml each were spread plated on selective agar respectively: MacConkey (Oxoid; targeting Gram-negative bacteria including *Escherichia* and *Klebsiella*), De Man-Rogosa-Sharpe (MRS; Difco) with 50 mg/L mupirocin (Oxoid; targeting *Bifidobacterium*), Baird-Parker (Oxoid; targeting *Staphylococcus*) and Slanetz and Bartley (Oxoid; targeting *Enterococcus*). Agar plates were then incubated both aerobically (MacConkey, Baird-Parke, and Slanetz and Bartley) and anaerobically (MRS agar only) at 37°C for 3 days. Next, 5 colonies from each agar plate were picked and were re-streaked for 3 consecutive times onto newly prepared agar plates to obtain pure isolates.

Bacterial isolates were cultured in appropriate media for overnight prior to genomic DNA extraction. DNA extraction was performed using the phenol-chloroform extraction method as described previously^58^. Briefly, PBS-washed bacterial cell pellets were resuspended in 2 ml of 25% sucrose in 10 mM Tris and 1 mM EDTA at pH 8.0, followed by enzymatic lysing step using 50 μl of lysozyme at 100 mg ml^-1^ (Roche). Next, 100 μl of Proteinase K at 20 mg ml^-1^ (Roche), 30 μl of RNase A at 10 mg ml^-1^ (Roche), 400 μl of 0.5 M EDTA at pH 8.0 and 250 μl of 10% Sarkosyl NL30 (Thermo Fisher Scientific) were added sequentially into the lysed suspension. The suspension was then subjected to an 1 h incubation on ice and then a 50°C water bath for overnight. On the next day, 3 rounds of phenol-chloroform-isoamyl alcohol (Merck) extraction were performed using 15 ml gel-lock tubes (QIAGEN), followed by chloroform-isoamyl alcohol (Merck) extraction prior to ethanol precipitation step and 70% ethanol cell wash for 10 min (twice). The DNA pellets were then air dried overnight, re- suspended in 300 μl of sterile pure water and stored in -80°C freezer prior to further analyses.

### Bacterial pure isolate Whole Genome Sequencing (WGS) and genome assembly

WGS of each pure bacterial isolate was performed on Illumina NextSeq 500 at the Quadram Institute (Norwich, UK) to generate 151-bp paired-end reads. Raw sequence reads (FASTQ) were firstly quality filtered (-q 20) with fastp v0.20.0^59^ before *de novo* genome assembly via genome assembler optimiser Unicycler v0.4.9b^60^ at default parameters that involved *de novo* genome assembler SPAdes v3.11.1^61^, also Bowtie2 v2.3.4.1^62^, SAMtools v1.7^63^ and sequence polishing software Pilon v1.22^64^. Contigs with lengths <500 bp were discarded from each draft genome assembly before subsequent analysis. All draft genome assemblies underwent sequence contamination check via CheckM v1.1.3^65^, and contaminated (sequence contamination >5%) and/or incomplete (genome completeness <90%) genome assemblies were excluded from further analysis (n=1) resulting in a total of 89 high quality pure isolate genomes subject to subsequent genome analyses. Taxonomic assignment (species level identification) was performed using gtdb-tk v1.5.1^66^. Genome assembly statistics were generated via sequence-stats v1.0^67^ and genome performed using Prokka v1.14.6^68^ (Supplementary Table 3).

### Recovery of Metagenome-Assembled Genomes

Shotgun metagenome raw reads (FASTQ) were firstly trimmed, adaptor-removed and quality-filtered using fastp v0.20.0^59^ (-q 20). Subsequently, host-associated sequences were removed via KneadData v0.10.0 with human genome (GRCh38.p13 - no ALT) bowtie2 index file retrieved from https://benlangmead.github.io/aws-indexes/bowtie (with options --bypass-trim and –reorder) to generate clean FASTQ reads that consist of purely bacterial sequences. These metagenome reads were then co-assembled with MEGAHIT v1.2.9^69^ prior to the reconstruction of Metagenome-Assembled Genomes (MAGs). Next, MetaWRAP v1.3.2^70^ was utilised to extract MAGs based on metagenome co-assemblies generated and metagenome clean reads via three binning software included in the pipeline MetaBAT v2.12.1^71^, MaxBin v2.2.6^72^ and CONCOCT v1.1.0^73^ using sub-module *metawrap binning*.

MAGs were then refined using sub-module *metawrap bin_refinement* to select the high-quality bins (MAGs) from each sample with completeness >90% and contamination <5% via checkm v1.1.3^65^. All MAGs were taxonomically ranked using gtdb-tk v1.5.1^66^ via module *gtdbtk classify_wf*. A total of 411 high-quality (completeness >90%, contamination <5%) MAGs (n=322) and isolate genomes (n=89) were recovered for subsequent analyses. Next, all MAGs and isolate genomes were dereplicated using dRep v3.2.2^74^ at ANI 99.9% as the strain-level inference cut-off (n=195).

### Taxonomic profiling for faecal metagenomes

After quality filtering steps as described above, briefly, shotgun metagenome raw reads (FASTQ) were quality-filtered using fastp v0.20.0^59^ (-q 20) and host-associated sequences were removed using KneadData; Kraken v2.1.2^75^ was utilised for taxonomic assignment for purified metagenome reads (Kraken2 standard Refseq indexes retrieved from https://benlangmead.github.io/aws-indexes/k2, May 2021), with confidence level set at 0.1. Bracken v2.6.2^76^ was then deployed to re-estimate the relative abundance of taxa at both genus and species level (-t set at 10 as recommended to reduce false positive) from Kraken2 outputs as recommended. To simplify data for visualisation purpose, minimal genera with <1000 reads across all samples were removed prior to further analysis (with average 520,000 resultant genera reads across all samples). Minor genera with <2% (relative abundance) across all samples were classified as ‘Others’ to simplify representations prior to data visualisation in R^77^ using library ggplot2^78^.

### Phylogenetic relatedness estimation

The Mash-distance sequence tree consisted of 411 genomes (including MAGs) was generated using Mashtree v1.2.0^79^ with 100 bootstrap replicates and option --mindepth 0, as in other cross-genus/species trees in this study. Distance tree was then mid-point rooted and visualised in iTOL v6^80^.

### Metagenomic functional profiling

Filtered shotgun metagenomic raw reads as described in previous method section (using fastp v0.20.0) were concatenated into a single FASTQ file per sample prior to running Humann v3.0.0 and Metaphlan v3.0.13 based on CHOCOPhlAn_201901 metaphlan database (built using bowtie2) retrieved from https://zenodo.org/record/3957592#.YSZDVS1Q3_o. Path abundance output files were then normalised and used for visualisation of microbiome functionalities.

*Antibiotic resistance genes, Multi-Locus Sequencing Typing (MLST) and plasmid replicons* Antibiotic resistance genes (ARGs) sequence search on all sequence data was performed via ABRicate v1.0.1 with options --minid=95 and --mincov=90 based on nucleotide sequence database ARG-ANNOT^81^ NT v6 (retrieved from https://www.mediterranee-infection.com/acces-ressources/base-de-donnees/arg-annot-2/), individually validated against ResFinder v4.0^82^ (only acquired ARGs) databases. The resistome analysis of infant gut metagenome was carried out via sequence search on metagenome co-assemblies of each sample, whilst strain-level resistome analysis was performed on individual genome assemblies. Antibiotic classes were sorted using outputs from ABRicate v1.0.1 based on antibiotic classes specified in ARG-ANNOT. Multidrug resistance efflux pumps were predicted using METABOLIC v4.0^83^.

MLST was predicted via mlst v2.19.0^84^ at default parameters through scanning contig files against the PubMLST^85^ typing schemes (https://pubmlst.org/) sited at the University of Oxford.

Plasmid replicons were predicted using ABRicate v1.0.1^86^ via PlasmidFinder sequence database v2^87^ with options --minid=90 and --mincov=90 on individual strain-level genome assemblies. Plasmid replicons were reclassified in two key groups - *inc* families and *rep* groups, respectively prior to visualisation using iTOL v6^80^.

### Klebsiella surface antigen typing

Kleborate v2.3.2^88^ was invoked with the –all flag to type the capsule (K) and LPS (O) antigen loci, using default parameters.

### Horizontal gene transfer analysis

Waafle v1.0^89^ was used with default parameters to find incidences of horizontal gene transfer in the dereplicated MAGs and whole genome sequences. To determine statistical significance between the antibiotic positive (n=237) and antibiotic negative (n=174) cohorts, a random permutation test was conducted to normalise the sample sizes. Here, the number of LGT events predicted by Waafle from either population were randomly sub-sampled 100 times, and these values were then subject to a Mann Whitney statistical test, and averaged, using Graphpad Prism v10.

### Enterococcus-associated plasmid transfer study via ex vivo colon model

*Enterococcus faecium* ARMA59 and ARMA73 were isolated from preterm infant stool samples that were part of the Baby-Associated Microbiota of the Intestine (BAMBI) study. Isolates were selectively cultured from stool samples using Slanetz and Bartley Medium (Oxoid), on which *Enterococcus* forms dark red colonies. Colonies were then streaked on Brain Heart Infusion agar (Merck) before being sent for whole genome sequencing. *E. faecium* 64/3, a plasmid-free derivative exhibiting high-level resistance to both rifampicin and fusidic acid, was utilized as the recipient strain for conjugation assays. This strain was originally isolated from a stool sample of a hospital patient^90^.

*Enterococcus faecium* ARMA59 and ARMA73 were used as donor strains in the *ex vivo* gut model experiment, while *Enterococcus* 64/3 was used as a recipient strain. Both ARMA59 and ARMA73 were confirmed for their gentamicin resistance (>130µg/ml) prior to the key experiment, while recipient strains 64/3 was phenotypically resistant to rifampicin (>25 µg/ml) and fusidic acid (>25 µg/ml).

Next, faecal slurry was prepared using *Enterococcus*-negative infant stool samples determined by selective agar plates (5 infants from PEARL study). Stool samples (1,965 mg) were mixed with 7 ml of reduced PBS to constitute the faecal slurry for the colon model experiment. Colon media were formulated based on our previous study with added vitamin solutions^91^. Briefly, 6 different media, M1 (500 ml), M2 (100 ml), M3 (100 ml), M4 (100 ml), M5 (200 ml), M6 (200 ml) were mixed aseptically in a sterile container. In addition, 200 ml of milk and vitamin mix (50 ml; pantothenate 10 mg/L, nicotinamide 5mg/L, thiamine 4mg/L, biotin 2mg/L, vitamin B12 0.5mg/L, menadione, 1mg/L and p-aminobenzoic acid 5mg/L) were also added, making the total volume 2000ml, and this was adjusted to a maximum 700 ml final total volume (for each of the media or additives). These colon media were then pH-adjusted to pH6.8.

For bacterial strain preparation, frozen bacterial stocks were plated on Brain Heart Infusion (BHI) agar plates for 24 h anaerobically at 37°C, followed by inoculating single colonies in 5ml BHI broth for overnight (16h) anaerobically at 37°C. Next, the cultures were homogenised and mixed (200 µl) with 10 ml of colon media (equivalent to 1:50 dilution), incubated at 37°C anaerobically overnight. The cultures were next washed (PBS + 3% cysteine) twice and pelleted.

Prior to experiments, faecal slurry was determined to be *Enterococcus*-free by both culturing (on Slanetz and Bartley Agar) and Whole Genome Sequencing approaches. DNA extraction of faecal slurry was performed via FastDNA Spin Kit for Soil (MP Biomedicals). Genomic DNA was then subjected to Whole Genome Sequencing via Illumina NovaSeq X to generate 151-bp paired-end reads. Raw sequencing reads (FASTQ) was processed as mentioned in previous section on metagenomic sequence analysis. Briefly, raw sequences were firstly quality-filtered with fastp v0.20.0 prior to taxonomic classification of sequence reads using Kraken v2.1.2^75^ (Kraken2 standardDB; --confidence 0.1) and further classified by Bracken v2.8^76^ (-t 10).

To set up the *ex vivo* colon model, the micro-Matrix fermentation system (Applikon Biotechnology) was used to model the human distal colon^92^. This system utilised a 24-well plate for batch fermentation. Four conditions were set up in the 24-well plate: 1) no donor, no recipient (control), 2) donor, no recipient, 3) no donor, recipient only, 4) donor and recipient strains. We have used a total volume of 5 ml in each well (slurry 26.6µl, donor strain inoculum 100µl, recipient strain inoculum 200µl, colon media negative control 100/200µl depending, colon media 4.67ml). The basic parameters of the colon model was set at pH6.8, temperature 37°C, and duration of experiment at 24 h. Samples were collected at 0h and 24h.

After the experiments, samples from 4 conditions then were re-streaked on Slanetz-Bartley agar plates supplemented or not with appropriate antibiotics: 1) no antibiotics, 2) 130 µg/ml gentamicin, 3) 25 µg/ml rifampicin + 25 µg/ml fusidic acid, 4) 130 µg/ml gentamicin + 25 µg/ml rifampicin + 25 µg/ml fusidic acid, to allow the selection of donor and recipient strains, and transformants. After 48h anaerobic incubation at 37°C, pure colonies were selected and cultured in media broth to obtain bacterial pellets sufficient for DNA extraction. Genomic DNA extraction of each isolate was performed via FastDNA Spin Kit for Soil (MP Biomedicals) as per manufacturer’s instructions. Well contents from 0h were also sampled and DNA extracted with FastDNA Spin Kit for Soil (MP Biomedicals) for shotgun metagenome sequencing to confirm the absence of *Enterococcus*.

Genomic DNA was whole-genome sequenced on the Nanopore MinION platform (R10.4.1 flowcell) to generate long-read raw sequences with basecalling via Dorado^93^. Raw sequences were filtered using filtlong v0.2.1^94^ to remove reads with <1000 bp. Filtered reads were then assembled via Flye v2.9^95^ (--nano-hq –scaffold -g 3.6m). Genome assemblies were subsequently polished using Medaka v1.11.3^96^ (*medaka_consensus* -m r1041_e82_400bps_sup_v4.3.0) to generate high-quality genome assemblies for subsequent analyses.

Isolate DNA was also whole-genome sequenced on Illumina NextSeq 2000 to generate 151- bp paired-end short reads (FASTQ) in parallel with long-read sequencing as described in previous paragraph. Raw reads were firstly trimmed with fastp v0.20.0^59^ (-q 20) prior to de novo genome assembly via genome assembler optimiser Unicycler v0.5.0^60^ at default parameters to generate draft genome assemblies for further analyses.

### Average Nucleotide Identity (ANI) estimation and bacterial strain-level identification

ANI was computed using fastANI v1.34^97^ at default parameters to compare genomes. Strain-level ANI cut-off was set at 99.9%, species-level cut-off was 95% (species boundary). To classify bacterial genomes at strain-level, dRep v3.2.2^74^ (*dRep dereplicate -- ignoreGenomeQuality*) was used at 99.9% ANI (-sa 0.999) to separate highly similar genomes.

### Data visualisation

Various bar plots, box plots, dot plots, pie charts and scatter plots were graphed via R v4.1.2^77^, using R libraries tidyverse v1.3.1^98^, ggplot2 v3.3.5^78^ and ggpubr v0.6.0^99^. R library vegan v2.6.2^100^ was used to construct non-metric multidimensional scaling (NMDS) plot for gut microbiome (taxonomic/functional profiling) data. Genomic region of contigs/gene comparison was performed via R library genoplotr v0.8.11^101^ or GenoFig v1.1.1^102^.

## Statistics

Statistical tests were performed via R base packages stats v4.1.2^77^, including Wilcoxon’s test and Shapiro–Wilk normality test which was used to formally test for data normality where appropriate. Spearman correlation tests were carried out via R library ggpubr v0.6.0^99^ function ggscatterhist. LEfSe^103^ was used to perform linear discriminant analysis (LDA) on functional profiling output data from Humann3^104^. Shannon diversity index of metagenome data was estimated via R library vegan v2.6.2^100^. Genome statistics were generated via sequence-stats v1.0^67^. R library dplyr v1.0.2^105^ was used frequently for data handling. Cohort characteristics were compared using either Wilcoxon test, Student’s t-test or Fisher’s exact test where appropriate, via R packages rstatix v0.6.0^106^.

## Data and code availability

Infant faecal sample metagenome sequencing raw reads are publicly available in the NCBI Sequence Read Archive (SRA) under accession no. PRJNA1191223. Sequencing raw reads and draft genome assemblies for 89 isolates generated in the present study are made available in the NCBI SRA and GenBank respectively, under accession no. PRJNA119225.

Sequencing raw reads and draft genome assemblies from long-read WGS on *Enterococcus* plasmid transfer study are publicly available in SRA (for raw sequencing reads) and GenBank (genome assemblies) respectively under accession no. PRJNA119226. All 322 metagenome-assembled genomes recovered from gut metagenome sample in this study and associated R scripts (data visualisation and statistical analysis) are available and shared via GitHub repository: https://github.com/raymondkiu/Infant-Resistome-Study

## Acknowledgements

This research was supported in part by the NBI Research Computing through the provision of a high-performance computing cluster. We would also like to thank the DNA sequencing team at Wellcome Trust Sanger Institute for performing genomic DNA sequencing for this study. This work was funded via a Wellcome Trust Investigator Award to L.J.H. (100974/C/13/Z) and support of the BBSRC Norwich Research Park Bioscience Doctoral Training Grant (BB/M011216/1; supervisor, L.J.H.; student, C.A.G.), Institute Strategic Programme (ISP) grant for Gut Health and Food Safety, BB/J004529/1 (L.J.H.), and ISP grant for Gut Microbes and Health BB/R012490/1 and its constituent project(s), BBS/E/F/000PR10353 and BBS/E/F/000PR10355 to L.J.H.; L.E.L., A.A.G., W.v.S and L.J.H were also supported by BBSRC grant BB/S017941/1. Work at Imperial Healthcare NICUs was supported by a programme grant from the Winnicott Foundation to J.S.K. and the National Institute for Health Research (NIHR) Biomedical Research Centre based at Imperial Healthcare NHS Trust and Imperial College London. K.S. was funded by an NIHR Doctoral Research Fellowship (NIHR-DRF-2011-04-128). We sincerely thank all clinical nurses at NNUH and Imperial Healthcare NICUs for collecting stool samples. We would like to give a special mention to research nurses Karen Few, Hayley Aylmer, and Zoe McClure for obtaining consent from parents and collecting samples.

## Author contributions

R.K., C.A. and L.J.H. conceived the study. R.K., A.A.G., L.E.L. and C.A. provided the methodology. R.K. and E.M.D. provided the software. L.J.H. and E.M.D. validated the study. R.K., E.M.D., A.A.G., L.E.L., A.C. and C.A. did the formal analysis. R.K., E.M.D., A.A.G., C.A. and A.C. carried out the investigations. K.S., L.E.L., A.G.S., P.C. and J.S.K. provided the resources. R.K., C.A. and K.S. curated the data. R.K. and L.J.H. wrote the original draft of the manuscript. W.vS., J.S.K. and L.J.H. supervised the study. R.K., A.A.G., W.vS., J.S.K., P.C. and L.J.H. reviewed and edited the manuscript. R.K. and C.A. administered the project. J.S.K. and L.J.H. acquired funds.

